# Decoding chromosome organization using CheC-PLS: chromosome conformation by proximity labeling and long-read sequencing

**DOI:** 10.1101/2024.05.31.596864

**Authors:** Kewei Xu, Yichen Zhang, James Baldwin-Brown, Thomas A. Sasani, Nitin Phadnis, Matthew P. Miller, Ofer Rog

## Abstract

Genomic approaches have provided detailed insight into chromosome architecture. However, commonly deployed techniques do not preserve connectivity-based information, leaving large-scale genome organization poorly characterized. Here, we developed CheC-PLS: a proximity-labeling technique that indelibly marks, and then decodes, protein-associated sites. CheC-PLS tethers dam methyltransferase to a protein of interest, followed by Nanopore sequencing to identify methylated bases - indicative of *in vivo* proximity - along reads >100kb. As proof-of-concept we analyzed, in budding yeast, a cohesin-based meiotic backbone that organizes chromatin into an array of loops. Our data recapitulates previously obtained association patterns, and, importantly, exposes variability between cells. Single read data reveals cohesin translocation on DNA and, by anchoring reads onto unique regions, we define the internal organization of the ribosomal DNA locus. Our versatile technique, which we also deployed on isolated nuclei with nanobodies, promises to illuminate diverse chromosomal processes by describing the *in vivo* conformations of single chromosomes.

## Introduction

Our understanding of chromosome organization has advanced significantly over the past few decades through the widespread application of genomic approaches. Chromatin immunoprecipitation (ChIP) relies on crosslinking proteins to DNA to record *in vivo* proximity. The enriched sequences can then be detected by various approaches, such as massively parallel (i.e., Illumina™) sequencing, and then mapped to the genome (Gilmour and Lis 1984; Furey 2012). Hi-C and related chromosome conformation capture (3C) approaches rely on the ligation of genomic DNA that was crosslinked and digested *in situ*. The frequency of sequencing reads that span two different regions of the genome is used to estimate *in vivo* proximity (Sati and Cavalli 2017; Dekker et al. 2002). These genomic approaches have been widely deployed to describe the location of chromosomal loci relative to one another and the association patterns of regulatory factors and nuclear scaffolds.

While both classes of approaches provide high-resolution, genome-wide protein-DNA and DNA-DNA proximity information, they have crucial shortcomings. First, since genomic DNA is sheared or digested, long-range information is effectively erased. This means that it is challenging to deduce whether a chromosomal transaction (e.g., protein binding) affects other biological processes that occur far away on the same DNA molecule. Second, these techniques provide statistical averages of a large cell population, masking variation between different cells. Third, these approaches capture a snapshot of the genome, limiting our understanding of dynamic events such as sliding along chromatin. Fourth, the reliance on short-read sequencing makes it difficult to unambiguously map sequencing reads onto sequence repeats, leaving the organization of repetitive regions mostly unknown.

To overcome these limitations, novel techniques are needed. Such techniques should be able to record *in vivo* proximity while accounting for the contiguity of the chromosome and for the movements of DNA and proteins relative to one another. Ideally, such techniques could be applied genome-wide, including to repetitive regions, and preserve the underlying heterogeneity between different cells. Several recent developments have started to chip away at this challenge. These include single-cell ChIP-seq and Hi-C (Zhou, Zhang, and Ma 2021; Schwartzman and Tanay 2015), Pore-C, which concatenates ChIP fragments to derive connectivity-based information (Deshpande et al. 2022), and DiMeLo-seq, which uses proximity labelling *in situ* to derive nucleosome positioning information (Altemose et al. 2022). Nonetheless, we still lack a versatile, robust genomic technique that overcomes the limitations of short-read-based approaches.

Here, we have developed a novel technique designed for decoding *in vivo* associations along single DNA molecules, which we call CheC-PLS: <u>ch</u>romosom<u>e</u> <u>c</u>onformation by <u>p</u>roximity labeling and long-read <u>s</u>equencing (pronounced “check, please”). CheC-PLS utilizes a chromosomal protein tethered to a DNA methyltransferase, which modifies nearby DNA sequences ((Kind et al. 2015; van Steensel, Delrow, and Henikoff 2001); Fig. 1A). Methylated sites are identified through Nanopore sequencing, which threads ultra-long DNA molecules through a protein pore without shearing or amplification and can simultaneously detect sequence information and base modifications (Hook and Timp 2023; Simpson et al. 2017). CheC-PLS offers the potential to provide single-molecule description of chromosome organization and connectivity, e.g., whether binding to two distant sites occurs concurrently, is mutually exclusive, or happens independently.

**Figure 1.**
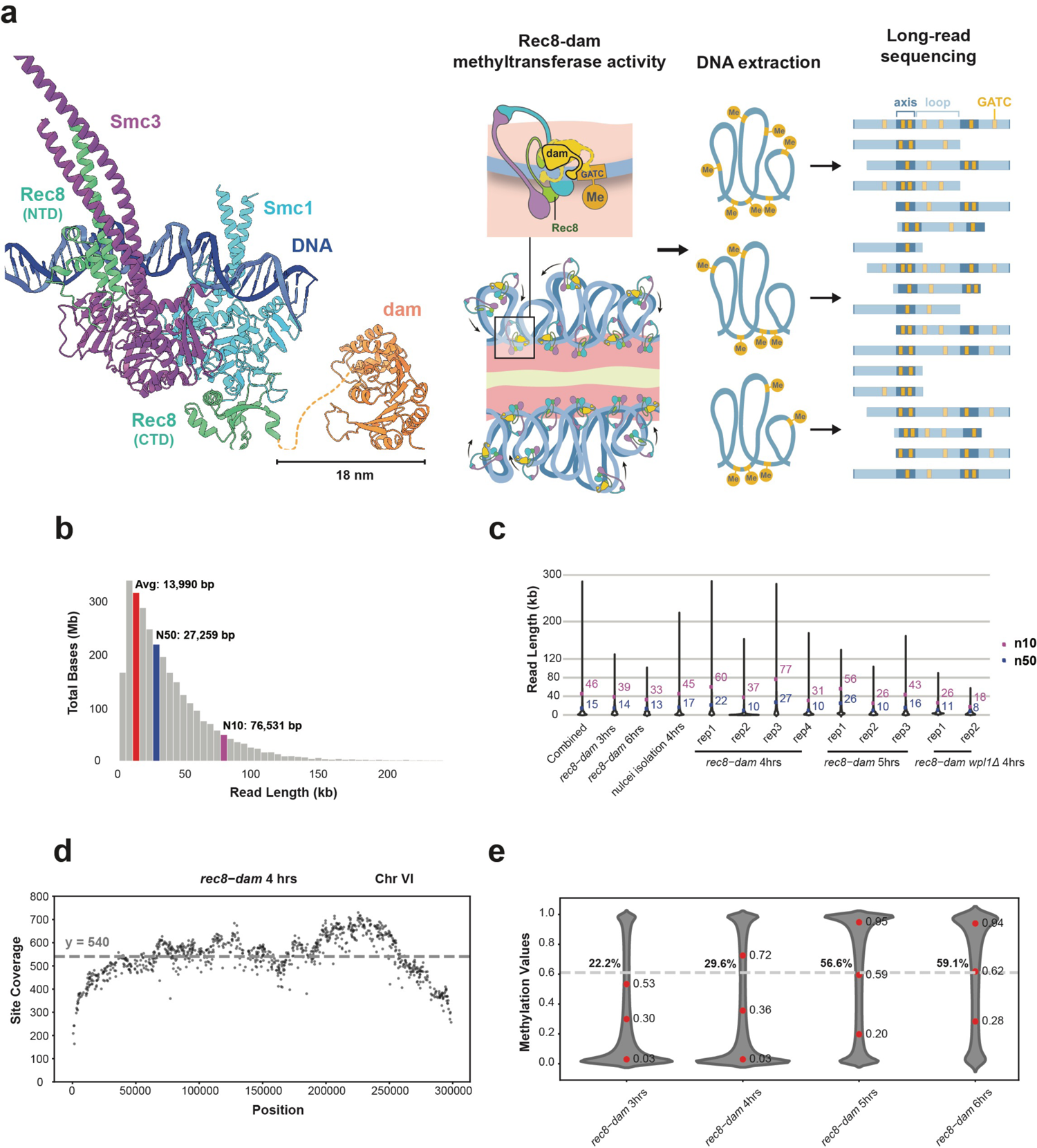
Validation of methylation calls on Nanopore sequencing reads. (a) Schematic representation of CheC-PLS. See text for details. Left, ColabFold projection image illustrating the association of Rec8-dam with Smc1, Smc3, and DNA molecules. The dashed yellow line represents the unstructured linker between Rec8 and dam. (b) Histogram of the total bases (read length * read numbers) binned by read length for a typical sequencing experiment. Red, average read length (14kb); blue, N50 read length (27kb); purple, N10 read length (77kb). (c) Violin plot depicting the distribution of read lengths across experiments, with N50 and N10 indicated in blue and purple, respectively. (d) Coverage of GATC sites along chromosome VI, 4 hours post-induction into meiosis. Each dot represents the number of reads that includes a particular GATC site. (e) Violin plots of all methylation values for each experiment. The fraction of reads with methylation values above 0.61 (considered to be methylated; indicated by a dashed lines). The red dots indicate values for 25th, 50th and 75th percentile.

As proof of CheC-PLS’ ability to provide novel insight into chromosome organization, we applied it to the meiotic chromosome axis - a conserved structure crucial for the successful production of gametes (Zickler and Kleckner 2023). The axis anchors the bases of chromatin loops, organizing them into a linear array (Fig. 1A). It is made of cohesins and other meiosis-specific structural proteins. (In budding yeast, the axis comprises the universal cohesin subunits Smc1 and Smc3, the meiosis-specific cohesin subunit Rec8, and the structural proteins Hop1 and Red1.) Cohesins are essential for the formation of the axes, where they contribute two key activities: topological entrapment of sister chromatids to mediate cohesion, and motor activity that extrudes chromatin loops through translocation along DNA (Sakuno and Hiraoka 2022; Yatskevich, Rhodes, and Nasmyth 2019).

Axis organization was first observed in electron micrographs of hypotonically-treated meiocytes, revealing chromatin loops emanating from rod-like structures (Rattner, Goldsmith, and Hamkalo 1981; Nebel and Coulon 1962). ChIP-based approaches revealed that axis proteins preferentially localize to distinct sites (‘peaks’) that are the base of chromatin loops (Blat et al. 2002; Panizza et al. 2011). ChIP also revealed that axis components can relocate as a consequence of transcription (Sun et al. 2015). Hi-C confirmed that the base of adjacent loops - the peaks in ChIP profiles of axis components - are in physical proximity (Schalbetter et al. 2019). Nonetheless, the details of the dynamic association of cohesin with chromosomes are poorly understood, as is whether binding to different axis-associated sites along the chromosome is coordinated.

## Results

### Rec8-dam is a functional axis protein

We developed CheC-PLS in budding yeast, a model organism devoid of significant endogenous DNA methylation (Hattman et al. 1978) and conducive to the efficient and synchronous induction of meiosis (Brar et al. 2012; Carlile and Amon 2008). We first sought to generate a functional methyltransferase fusion protein. Most of our attempts to attach various methyltransferases to meiotic axis components resulted in spore viability defects consistent with defective axis formation (Supplementary Fig. 1a). Nevertheless, we generated an endogenously-tagged, functional construct, Rec8-dam. Rec8 is the meiosis-specific kleisin subunit of cohesin (Klein et al. 1999; Watanabe and Nurse 1999) and dam is a bacterial DNA methyltransferase that methylates adenine in the context of a GATC sequence (Geier and Modrich 1979). Upon induction into meiosis, homozygous Rec8-dam cells sporulated at similar rates to controls (80% and 76%, not significant) and displayed only slightly lower number of viable spores per ascus (3.8 and 3.1, p < 0.05), suggesting the transgene does not dramatically compromise the functionality of Rec8, whose function is required for accurate meiotic chromosome segregation (Supplementary Fig. 1b).

### Long-read sequencing using Nanopore

CheC-PLS requires ultra-long sequencing reads that could capture long-range regulation on the same DNA molecule. We extracted high-molecular-weight genomic DNA from budding yeast meiocytes by adapting a lysis protocol (Erwan Denis, Sophie Sanchez, Barbara Mairey, Odette Beluche, Corinne Cruaud, Arnaud Lemainque, Patrick Wincker, Valérie Barbe 2018). Briefly, the cell wall was removed by zymolase to form spheroplasts, which were then gently lysed. Following protease and RNase digestion, masses of precipitated DNA were ‘fished’ with a pipette, washed and rehydrated. Pulsed-field gel electrophoresis revealed that the average length of DNA molecules in our preparations exceeded 200 kb (Supplementary Fig. 1c).

Nanopore MinION sequencing yielded reads with an average N50 of 15kb and N50 in specific experiments around 30 kb (Fig. 1c). N10 (the length of the top 10% of reads) averaged 46kb, and the longest sequencing reads exceeded 300 kb, surpassing the length of some budding yeast chromosomes (Fig. 1c). We obtained an average of 2,400Mb per experiment, with the majority of the budding yeast genome boasting >200-fold coverage (Fig. 1d).

### Methylation detection

Following initial base-calling and alignment to the budding yeast genome (Genome assembly ASM205788v1; base-calling by guppy; (J.-X. Yue et al. 2017)), we detected adenine methylation using Remora. Remora assigns each GATC site a methylation value ranging from 0 to 1, with higher values indicating a greater likelihood of methylation. Using *E. coli* DNA that is either completely methylated or lacks methylation altogether (*dam^+^ dcm^+^* and *dam^-^ dcm^-^*, respectively), we identified 0.61 as a threshold that yields the lowest false identification rate, <16% (Supplementary Fig. 1f). (Future applications could improve accuracy at the expense of resolution by smoothing the data, which reduces the error rate to 8.3% [Supplementary Fig. 1f].)

To examine potential sequence biases in methylation patterns or in methylation calling, we analyzed genome wide methylation patterns of naked budding yeast DNA methylated with recombinant dam *in vitro*. Our analysis revealed that methylation was distributed mostly evenly along the chromosome, with the exception of several troughs likely attributable to low coverage (Supplementary Fig. 2e).

### Aggregated CheC-PLS data recapitulates Rec8 ChIP profile

Rec8 is expressed and loads onto chromosomes at meiotic S-phase, which occurs at around 3 hours after induction into meiosis, and remains chromosome-associated throughout meiotic prophase (Klein et al. 1999). To investigate the binding pattern of Rec8, we synchronously induced meiosis in *rec8-dam* homozygous strains and collected cells after 3, 4, 5 and 6 hours. The latter two timepoints were analyzed in an *ndt80* deleted strain (*rec8-dam ndt80Δ*), where cells are arrested at the pachytene sub-stage of meiotic prophase with fully assembled axes.

Since budding yeast does not harbor any adenine demethylases (Fedeles et al. 2015), methylated sites are not diluted by DNA replication and are expected to accumulate throughout meiosis. This was indeed the case. The fraction of methylated sites increased with time in meiosis: at 3 hours 22.2% of GATC sites were methylated, and this number increased to 29.6% and 56.6% at 4 and 5 hours after meiotic induction, respectively (Fig. 1e). We observed no further increase between 5 and 6 hours (56.6% and 59.1% at 5 and 6 hours, respectively), likely due to the lack of available unmethylated adenines.

When it first loads onto chromosomes, Rec8 is enriched at the centromeric regions (Klein et al. 1999; Sun et al. 2015). Consistent with this preference, methylation accumulated at the ∼10 kb surrounding the centromeric regions during the early stages of meiosis (3- and 4-hours post-induction; Fig. 2a,b). However, as meiosis progressed (at 5-hours post-induction), methylation outside of the centromeric regions became more prominent, reaching similar methylation levels as the centromeres (Fig. 2a,b).

**Figure 2.**
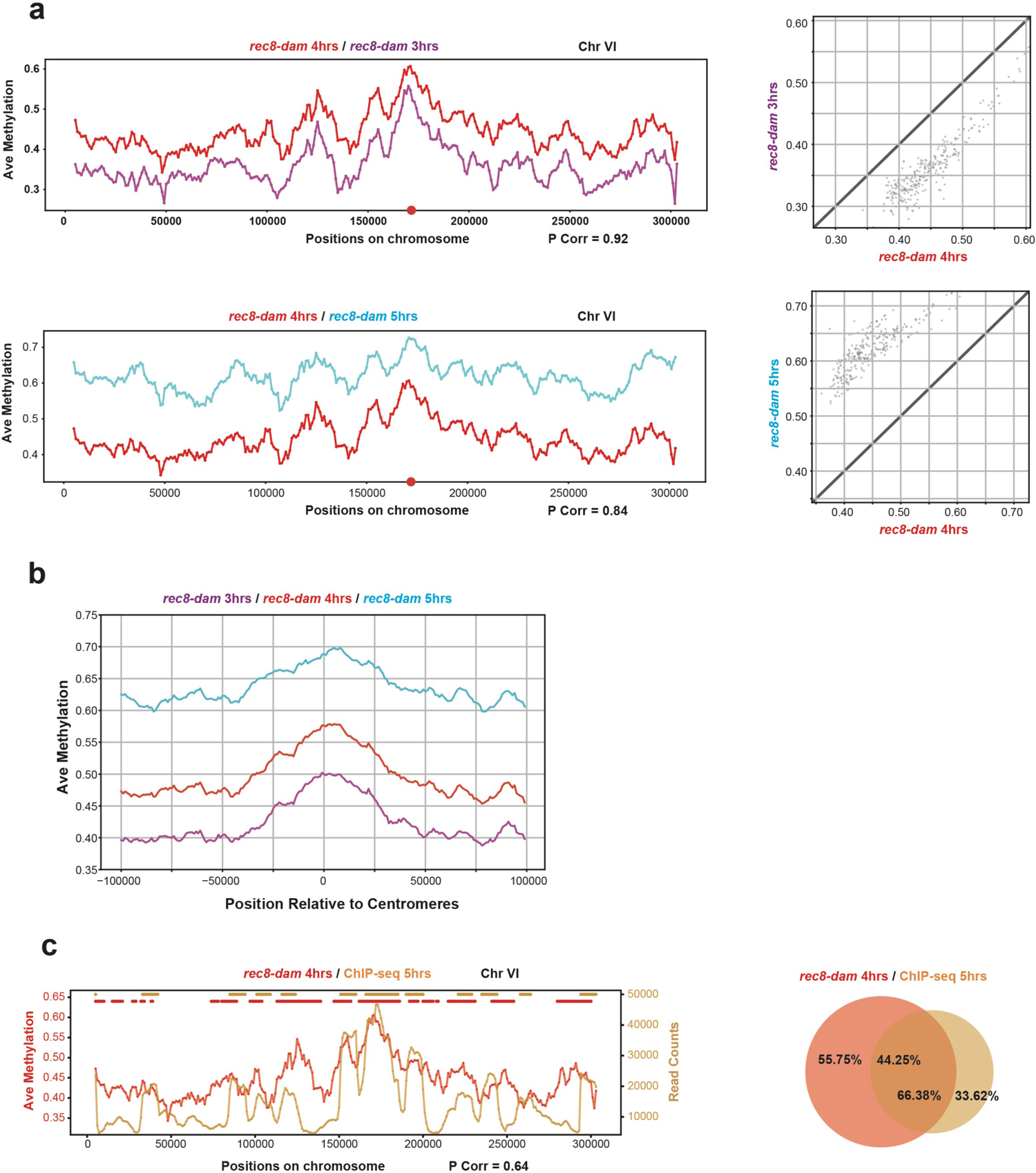
CheC-PLS data is robust and recapitulates ChIP-seq data (a) Left, averaged methylation across chromosome VI, with 3, 4 and 5 hours in meiosis represented by purple, red and cyan lines, respectively. The window size is 10 kb. The red dot indicates the centromere. Right, scatter plot between pairs of time points. Diagonal line indicates unchanged methylation. (b) Averaged methylation around the centromeres for all 16 chromosomes at 3, 4 and 5 hours (purple, red and cyan, respectively). The window size is 10 kb. See Supp. Fig. 2g for the complete data. (c) Left, averaged methylation across chromosome VI for CheC-PLS *rec8-dam* at 4 hours (red) and read count for Rec8 ChIP-seq at 5 hours (brown) after induction into meiosis. Red and brown dots above the plots indicate identified peaks. Pearson correlation between the datasets is 0.64. Right, overlap between peaks identified in the CheC-PLS and ChIP-seq datasets.

The methylation pattern throughout the genome exhibited a high correlation between biological replicates (cultures induced into meiosis in separate experiments and processed separately; Supplementary Fig. 2a) and between different timepoints (Fig. 2a). The Pearson correlation coefficient (P corr) was 0.91 between biological replicates, 0.92 between 3 and 4 hours, and 0.84 between 4 and 5 hours. Notably, there was no correlation between the CheC-PLS methylation pattern and sequencing depth or density of GATC sites (Supplementary Fig. 2d).

To further validate our observations, we compared the CheC-PLS signal to previously obtained ChIP-seq data (Fajish et al. 2024). It is important to note that ChIP-seq captures the association of proteins with DNA at the moment of crosslinking, whereas CheC-PLS records cumulative association. Despite these differences, Rec8 ChIP-seq and Rec8 ChIP-seq data revealed significant similarities (Fig. 2C, *left*; P corr = 0.64). This level of correlation is similar to that observed between ChIP-seq profiles of different axis components (Panizza et al. 2011). We also note the high degree of overlapping peaks in the signal of these two very different approaches (Fig. 2C, *right*).

The correlations between different CheC-PLS timepoints and between CheC-PLS and ChIP-seq validate the functionality of CheC-PLS, and, specifically, the robustness of proximity methylation *in vivo* and methylation calling of Nanopore reads by Remora. Our data also confirms that the identified methylation sites are genuine Rec8-associated sites, rather than biological artifacts or biases stemming from sequencing or methylation calling.

### Single-read analysis reveals long- and short-range coordination

So far, we have analyzed CheC-PLS reads in bulk. By averaging methylation values across many reads, we recapitulated the known genomic distribution of Rec8 (Fig. 2). The distinguishing feature of CheC-PLS, however, are sequencing reads that preserve the relationship between binding events along single, long molecules of DNA. Below, we harness this information to reveal how is Rec8’s association with meiotic chromatin is regulated over long distances.

Cursory analysis of reads hinted that methylation at nearby sites is correlated (Fig. 3a, asterisks). To systematically quantify such effects, we calculated the coefficient of coincidence, ln(CoC), between the methylation status of different GATC sites on the same read. ln(CoC)>0 indicates an increased likelihood of similar methylation status (either both methylated or both unmethylated), a situation known as positive interference. ln(CoC)<0 indicates that methylation on one site decreases the likelihood of methylation of nearby sites (negative interference), while ln(CoC)=0 indicates independent (random) methylation events.

**Figure 3.**
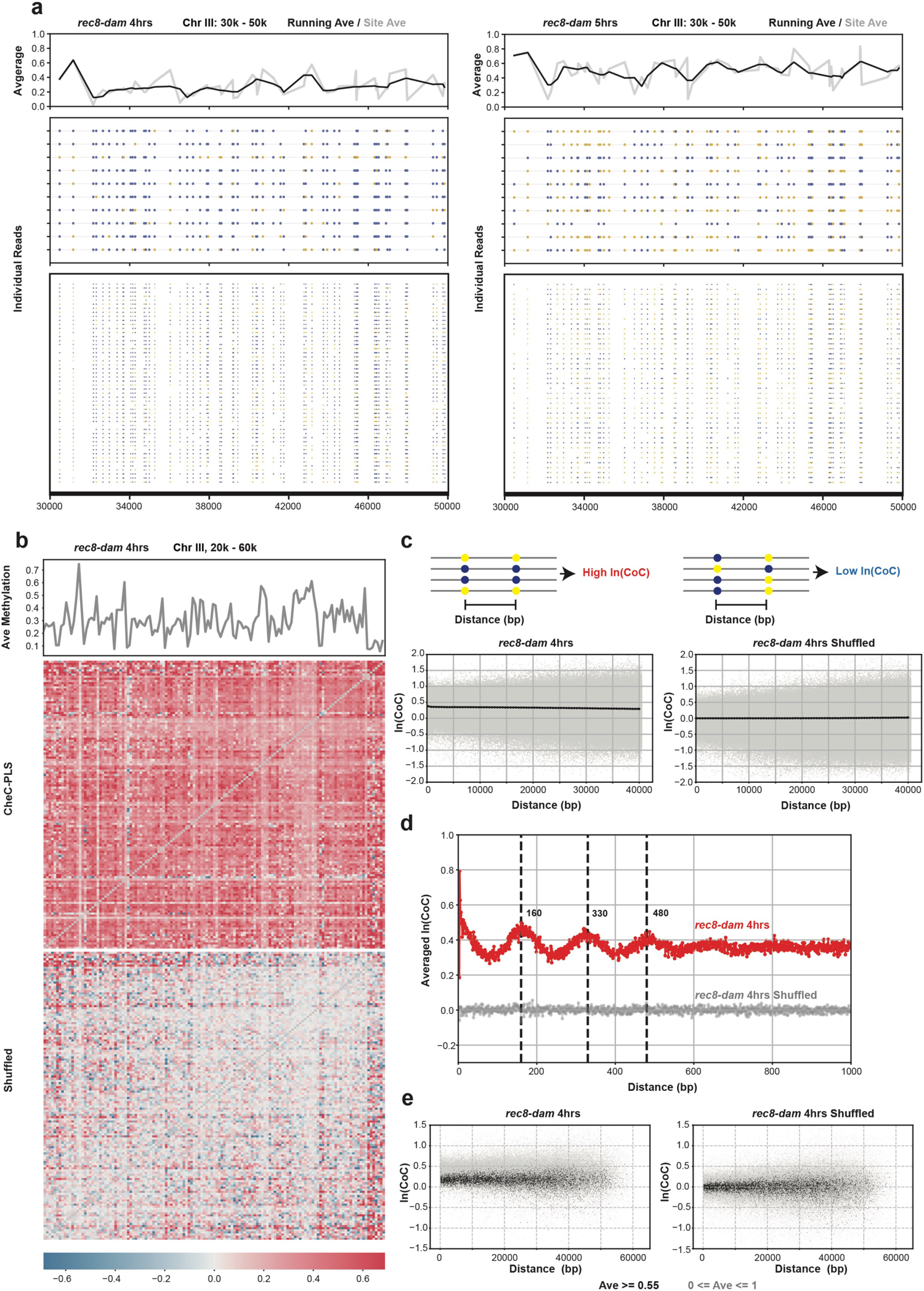
Short- and long-range correlation in methylation patterns on CheC-PLS reads (a) Single-read binding profile showing heterogeneity in Rec8 association along chromosome III, positions 30-50 kb, at 4 and 5 hours. Averaged methylation for each GATC site is plotted in grey, with a running average in black. Yellow dots indicate methylated GATC sites, and blue dots indicate unmethylated sites. (b) Heatmap of ln(CoC) for each pair of sites between 20 kb and 60 kb on chromosome III. ln(CoC) ranges from blue to red. Top, average methylation plot at each GATC site in this region. Bottom, the shuffled dataset eliminates the positive correlation. (c) Scatter plot of averaged ln(CoC) by distance between each pair of sites. Grey dots indicate each pair of sites, and black trend line indicate the binned average ln(CoC). Right, shuffled data showing average ln(CoC) close to 0, indicating uncorrelated events. Bin size = 500 bp. (d) Average ln(CoC) in the first 1kb (unbinned), with the 4 hours data in red and shuffled data in gray. Vertical dashed lines indicate the local maxima. (e) Sites with high methylation averages (≥ 0.55; black) exhibit lower ln(CoC). Right, shuffled data.

When analyzed over long ranges (GATC sites separated by 0.5-40 kb) we observed a moderately positive CoC, which declined slightly with increasing distance between methylated sites, from ln(CoC)=0.38 at 0.5 kb to ln(CoC)=0.29 at 40 kb (Fig. 3c, *left*). As a control, we generated datasets that retained the average methylation at each GATC site but shuffled methylation states between different reads (Fig. 3c, *right*; referred to as ‘shuffled’; see Methods). As predicted, ln(CoC)=0 in the shuffled datasets. We hypothesize that long-range coincidence is a result of the movement of Rec8 relative to the DNA molecules. This movement likely reflects loop extrusion by cohesin, although it might also be generated by sliding of cohesin rings on DNA, and/or unloading of cohesin followed by nearby reloading. We further test this idea using mutants and by analyzing isolated nuclei, below.

Cells in different meiotic timepoints exhibited a similar trend of minor decline with increasing distance between GATC sites, although the asymptotic value was lower at later time points (0.09 at 0.5 kb to 0.07 at 10 kb in the 5-hour data; Supplementary Fig. 3b, *left*). The lower ln(CoC) is likely due to the saturation of methylated GATC sites, diluting the effects of unique binding events and reducing our statistical power to detect coincidence. However, it may also reflect an underlying shift in the fraction of mobile cohesins or in the kinetics of cohesin translocation.

Over shorter distances (up to 1 kb) we observed a striking sinusoidal pattern, with peaks spaced ∼165bp apart and declining in amplitude with growing distance between GATC sites (Fig. 3d). We observed reduced amplitude (designated *û*, defined as the difference between the first trough and first peak) with increased time in meiosis (*û* = 0.12, 0.13 and 0.06 at 3, 4 and 5 hours), and ln(CoC)=0 for the shuffled datasets, similar to the patterns of long-range coincidence. The periodicity (∼165bp) is conspicuously similar to the predominant spacing between adjacent nucleosomes *in vivo* (Chereji et al. 2018). Notably, it is not merely a reflection of the spacing between GATC sites in the budding yeast genome (Supplementary Fig. 1d). Given that the predicted length of the protein linker that connects dam to Rec8 is roughly the same size as the diameter of a nucleosome (linker: 10nm; Fig. 1a; nucleosome = 11nm; (Luger et al. 1997)), we hypothesize that stacked adjacent nucleosomes face Rec8-dam, resulting in the higher coincidence of methylating sites that are ∼165bp apart.

To examine the genomic distribution of ln(CoC), we created heatmaps where GATC sites in the genome are placed on the x- and y-axes and each pixel represents ln(CoC) between a pair of GATC sites. Gray regions away from the diagonal represent pairs of GATC sites where not enough reads spanned both sites (Fig. 3B and Supplementary Fig. 3d). These heatmaps recapitulated our observations above: most pixels exhibited positive ln(CoC), and ln(CoC) did not dramatically decrease with increasing distance between GATC sites (represented in the heatmap as the distance from the diagonal). Most pixels in the shuffled datasets exhibited ln(CoC) close to 0.

Interestingly, we observed a weak inverse correlation between average methylation and ln(CoC), meaning that regions with high cohesin occupancy exhibited lower coincidence (Fig. 3e). A possible interpretation of this finding is that cohesin at the base of chromatin loops, which probably represents ‘cohesive’ cohesin that mediates sister-chromatid cohesion, is less mobile, resulting in lower ln(CoC). In contrast, the mobile, loop-extruding cohesins translocate along DNA and methylate GATC sites along the way, resulting in higher ln(CoC).

### Cohesin binding in the absence of Wpl1

Wpl1 is a conserved cohesin regulator that removes a subset of cohesin molecules from chromosomes. In its absence, cohesins accumulate on chromosomes, and, due to a smaller pool available for reloading, the potential for loop extrusion is reduced (Barton et al. 2022; Hong et al. 2019; Challa et al. 2016).

To test the potential effects of increased cohesin residency and decreased loop extrusion, we analyzed CheC-PLS data from homozygous *wpl1Δ rec8-dam* diploids undergoing meiosis. When compared to meiosis in the presence of *WPL1*, we noted very similar overall binding pattern (Fig. 4a; P corr = 0.95), as was previously reported for ChIP-seq profiles in *wpl1Δ* meiocytes (Barton et al. 2022; Hong et al. 2019; Challa et al. 2016). The similar methylation levels in cells with and without *WPL1* contrasts with the higher cohesion ChIP-seq signal in *wpl1Δ* cells (Barton et al. 2022), highlighting the contribution of cohesin dynamics to CheC-PLS signal. ln(CoC) values were lower in *wpl1Δ* cells over long-ranges, dropping from 0.38 in cells with *WPL1* at 0.5 kb, to 0.26 in *wpl1Δ* cells at 0.5 kb (Fig. 4b,d). The reduced coincidence was apparent despite the similar average methylation (29.6% versus 30.5% for *WPL1* and *wpl1Δ* cells; Supplementary Fig. 4b), suggesting the lower ln(CoC) is not a consequence of saturated methylation sites. Instead, our analysis suggests that ln(CoC) requires cohesin removal and re-loading, and suggests the observed coincidence is a consequence of cohesin movement on DNA. Consistent with this idea, the amplitude of the short-range, presumably nucleosomal, signal was also similar (*û* = 0.13 and 0.10 for cells with and without *WPL1*, respectively).

**Figure 4.**
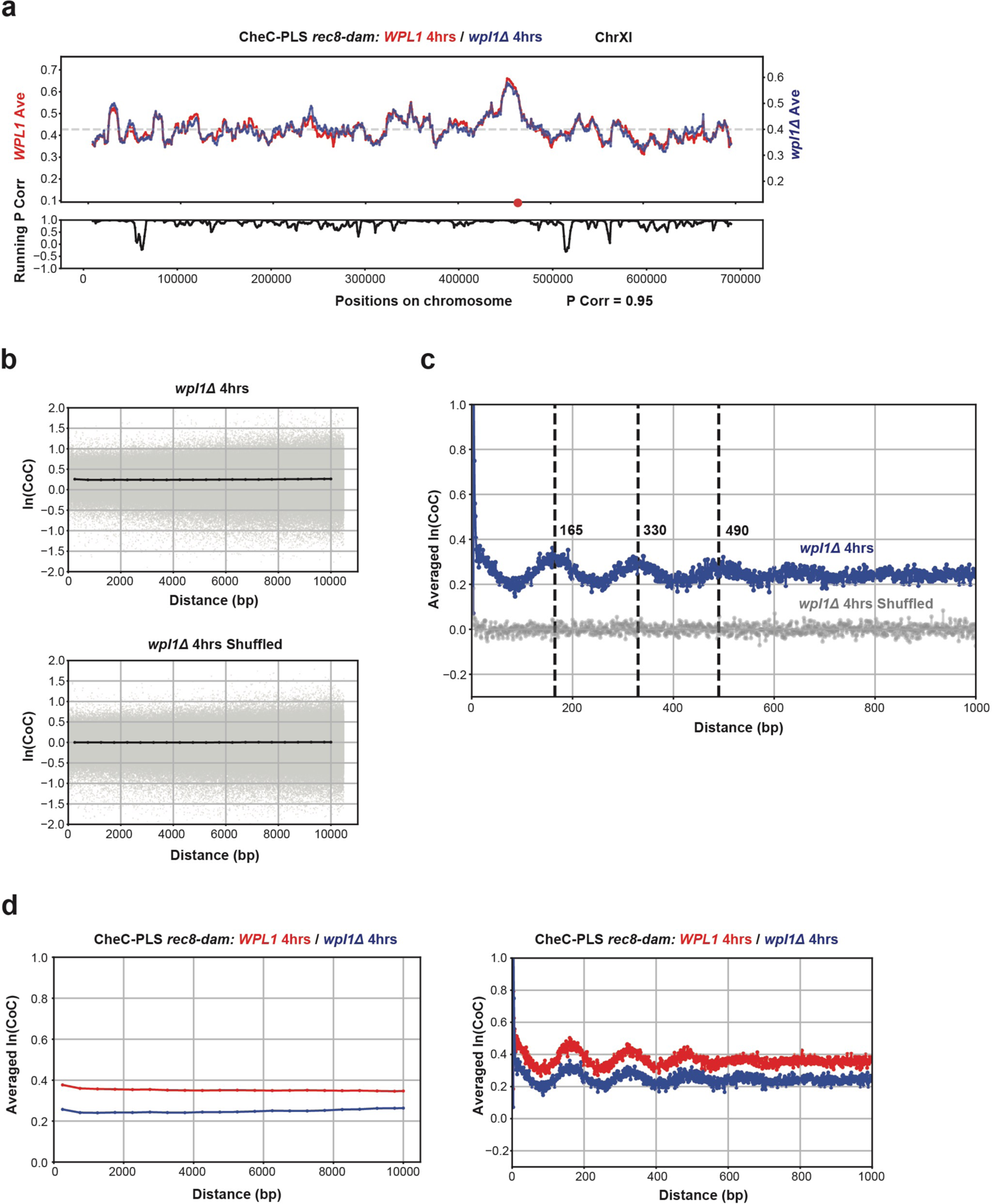
*wpl1* deletion does not alter Rec8 association patterns but reduces ln(CoC). (a) Average methylation plot for *WPL1* (red) and *wpl1Δ* (blue) at 4 hours on chromosome XI. Pearson correlation = 0.95. Window size = 10kb. Bottom, running Pearson correlation. (b) Averaged ln(CoC) by distance between pairs of sites, ranging from 0 to 10 kb, with a bin size of 500 bp. (c) Zoomed-in view of (b) with no binning. The *wpl1Δ* is shown in blue and the shuffled dataset in grey. (d) Comparison of averaged ln(CoC) by distance between *WPL1* (red) and *wpl1Δ* (blue). The analysis spans from 0 to 10 kb with a bin size of 500 bp (left) and from 0 to 1 kb without binning (right).

### CheC-PLS on purified meiotic nuclei

To further study the effects of cohesin dynamics, we wanted to deploy CheC-PLS in conditions that eliminate cohesin movement. Once removed from cells, nuclei are depleted of metabolites, grinding enzymatic processes to a halt. These processes include ATP-dependent translocation of cohesin along with other sources of both active and secondary chromosome movements.

To adapt CheC-PLS for *in situ* methylation, we isolated nuclei from yeast meiocytes expressing Rec8-GFP and incubated them with recombinant GFP-binding nanobodies fused to dam (GBP-dam), followed by DNA purification and processing as above (Fig. 5a; labelled ‘isolated nuclei’). This variant of CheC-PLS is conceptually analogous to other recently developed *in situ* approaches, such as DiMeLo-seq, nanoHiMe-seq and BIND&MODIFY (Altemose et al. 2022; W. Li et al. 2023; Weng et al. 2023).

**Figure 5.**
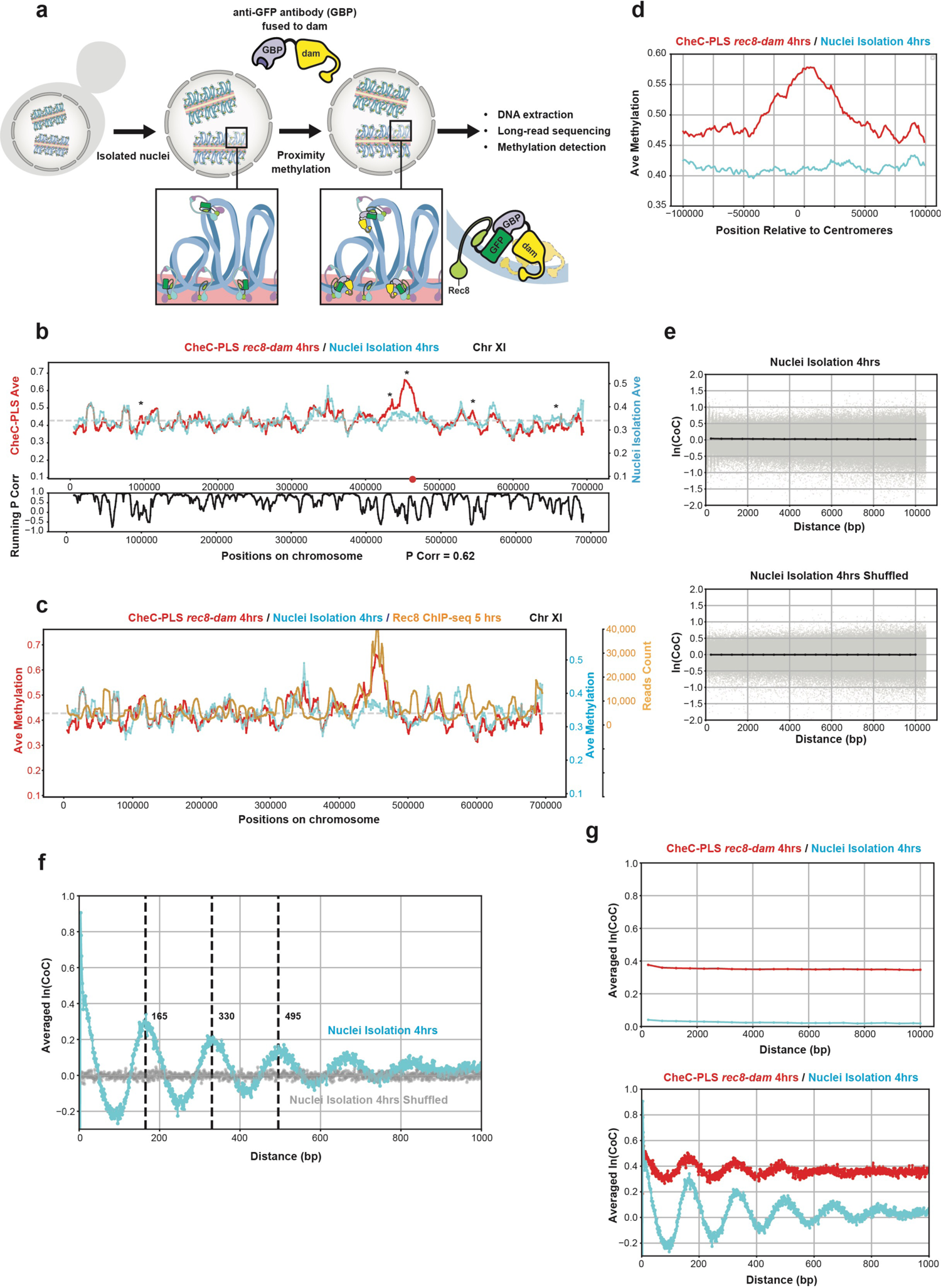
CheC-PLS on isolated nuclei (a) Schematic diagram illustrating deployment of CheC-PLS on isolated nuclei. See text for details. (b) Average methylation plots for CheC-PLS *rec8-dam* (red) and nuclei isolation (cyan), both at 4 hours on chromosome XI. Note lack of enrichment at the centromere (red dot) in the isolated nuclei. Window size = 10 kb. Bottom, running Pearson correlation. (c) Average methylation plots on chromosome XI for CheC-PLS *rec8-dam* at 4 hours (red), nuclei isolation at 4 hours (cyan), and Rec8 ChIP-seq at 5 hours (brown). (d) Averaged methylation around the centromeres for all 16 chromosomes for in vivo CheC-PLS *rec8-dam* (red) and isolated nuclei (cyan). The window size is 10 kb. See Supplementary Fig. 4a for complete data. (e) Average ln(CoC) by distance between sites; window size = 500bp. Note the very low ln(CoC) even at adjacent sites. (f) Zoomed-in view of the ln(CoC) plot in the first 1kb with no binning, showing more pronounced periodicity. (g) Comparison of averaged ln(CoC) by distance between CheC-PLS *rec8-dam* (red) and nuclei isolation (cyan) at 4 hours. The analysis spans from 0 to 10 kb with a bin size of 500 bp (top) and from 0 to 1 kb without binning (bottom).

The average methylation pattern in meiotic nuclei treated with GBP-dam was similar to *in vivo* CheC-PLS (Fig. 5b). However, there were also important differences. Some peaks that were present in the *in vivo* CheC-PLS data were missing in the nuclei data (Fig. 5b; asterisks: five peaks for chromosome XI). Some of these missing peaks may represent cohesin loading sites or other sites that are only occupied in earlier stages of meiosis. These regions will no longer be in proximity to cohesins in the isolated nuclei. One prominent class of cohesin peaks that were missing in the nuclei data were around the centromeres, where methylation was not enriched on any of the 16 chromosomes (Fig. 5d). The reason for the lack of centromeric signal is unclear, since centromeric DNA in enriched in ChIP-seq profiles of cohesins at the same meiotic stage (Fig. 5c; e.g., (Fajish et al. 2024)). A potential explanation for this depletion is a unique state of centromeric chromatin in native preparations (Krassovsky, Henikoff, and Henikoff 2012), which might affect the accessibility to GBP-dam.

When analyzed for the coincidence of methylation, CheC-PLS on isolated nuclei exhibited a distinct pattern. Over long distances, we observed an almost complete loss of coincidence, with ln(CoC) = 0.04 at 0.5 kb (Fig. 5e) - much lower than *in vivo* methylated meiocytes at the same timepoint (ln(CoC) = 0.38; Fig. 3c). This observation lends support to the idea that positive ln(CoC) reflects sliding of cohesins on chromatin, since sliding requires either active ATP hydrolysis (e.g., for transcription or loop extrusion) or indirect chromosome movements, which are both eliminated in isolated nuclei. Strikingly, the signature for short-range coincidence was dramatically increased in isolated nuclei, as indicated by higher amplitudes (*û* = 0.44 for isolated nuclei *versus* 0.13 for *in vivo* CheC-PLS; Fig. 5f,g). This observation is consistent with short-range coincidence resulting from stacked nucleosomes that are less mobile in isolated nuclei, where processes such as transcription and chromatin remodeling are not taking place.

### CheC-PLS reveals internal organization of the rDNA locus

Techniques like ChIP-seq and Hi-C rely on short sequencing reads, limiting the ability to uniquely map reads onto repetitive regions. Reads mapping to repeats are commonly excluded or pooled together, masking potential differences in binding patterns. The long reads used by CheC-PLS offer the potential to detect binding patterns and define genome organization in repetitive regions.

To test this ability, we focused on the ribosomal DNA (rDNA) locus in budding yeast, which harbors 100-200 tandem copies of a 9.1kb repeat that encode the RNA subunits of the ribosome (Salim and Gerton 2019). We mapped methylation sites along the rDNA array by anchoring them onto unique regions abutting the rDNA in a modified genome that included 20 rDNA repeats. (The reference budding yeast genome includes only two repeats; see Methods). This strategy gave us unprecedent view into cohesin association patterns in a native rDNA locus (Fig. 6a).

**Figure 6.**
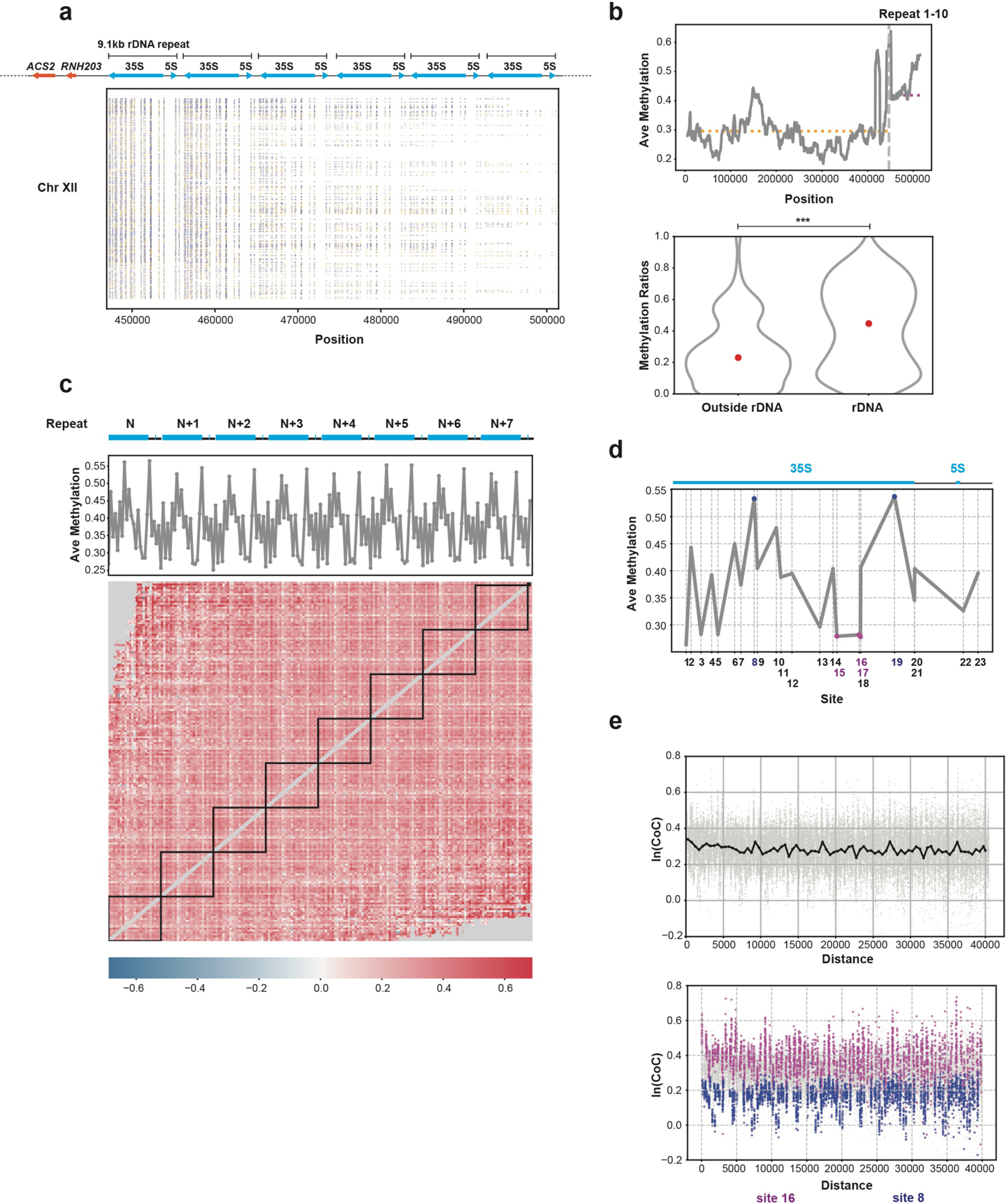
CheC-PLS define Rac8 association pattern in the rDNA region (a) Single-read methylation profile showing heterogeneity between single reads mapped to the rDNA region by anchoring it to the unique sequences to its left. Yellow dots indicate methylated GATC sites, and blue dots indicate unmethylated sites. The chromosome shown is chromosome XII, from 446 kb to 502 kb. Top, gene structure showing the 9.1kb rDNA repeat (blue) and the first two unique genes to the left of the rDNA locus (red). (b) Top, average plot showing Rec8 enrichment in the rDNA region compared to regions outside the rDNA. Dashed lines indicate average methylation outside (yellow) and inside (magenta) the rDNA array. Bottom, a violin plot showing the average methylation levels in 1000 randomly selected 9.1 kb windows outside the rDNA and averaged methylation levels for all rDNA repeats. (c) Heatmap of ln(CoC) in pooled reads of the rDNA region from repeat N to repeat N+7, showing inter- and intra-repeat correlations. (d) Binding patterns to the rDNA, averaged across all reads and across all repeats. Numbers along the x-axis indicate the 23 GATC sites in the 9.1kb repeat. (e) Top, ln(CoC) by distance in pooled reads mapped to the rDNA. A repetitive pattern is observed, matching the 9.1kb periodicity of the rDNA repeats. Bottom, sites with low methylation (site 16, magenta) are associated with high ln(CoC), while sites with high average methylation (site 8, blue) are associated with low ln(CoC).

When comparing average methylation patterns before and within rDNA repeats, we observed significant positional effects. The methylation signal was higher within the rDNA repeats compared with the region outside these repeats (Fig. 6b). This suggests that the rDNA region is highly organized during meiosis, a finding consistent with previous research (Vader et al. 2011). Within the rDNA, we did not observe a strong effect of proximity to the unique sequences outside the array either in the overall methylation levels or in ln(CoC) (Supplementary Fig. 5a).

We also stacked together all the reads containing rDNA sequences, independent of their position on the chromosome. The large number of very long reads in the rDNA array allowed us to test for potential coincidence that depends on the position of rDNA repeats relative to one another.

We found a significant level of coincidence between methylation of the rDNA repeats on the same sequencing reads, which was eliminated in the shuffled dataset (Fig. 6e, Supplementary Fig. 5b). Interestingly, ln(CoC) did not significantly decrease between repeats that are further apart (Fig. 6c; e.g., repeat N and N+3). As we observed above for unique sequences, ln(CoC) between rDNA repeats was consistently higher for low-methylated regions (Fig. 6e).

Each rDNA repeat contains 23 GATC sites, and we observed high Rec8 association at sites #8 and #19 and low association at sites #15, #16 and #17 (Fig. 6d). This methylation pattern is similar to the one observed for meiotic budding yeast by ChIP-seq, but differs from the ChIP-seq pattern of mitotic cohesins (Glynn et al. 2004; Costantino et al. 2020).

We also compared Rec8 rDNA profiles in the *wpl1Δ* meiocytes and in isolated nuclei. Patterns were very similar between cells with and without *WPL1*, including similar binding profile to each repeat, higher methylation within the rDNA array, and generally positive ln(CoC) that did not dramatically decrease with distance between the repeats (Supplementary Fig. 5c,d). Analysis of isolated nuclei yielded changes relative to the *in vivo* datasets consistent with the genome-wide differences. While local binding patterns were similar, inter-repeat ln(CoC) was completely eliminated, with only weak intra-repeat ln(CoC) signal remaining (Supplementary Fig. 5e,f). The absence of inter-repeat ln(CoC) signals is consistent with cohesin loop extrusion traversing multiple rDNA repeats. The intra-repeat positive signals suggests that cohesins organize them into distinct units. This result reiterates our conclusion that large-scale coordination in methylation status reflects cohesin dynamics on DNA.

## Discussion

The basic organizational unit of chromatin - the nucleosome - is well-characterized, as are some of its large-scale packaging principles, where cytological approaches have been extensively deployed. However, our mechanistic and functional understanding of intermediate scales - including chromosome loops, topologically associated domains (TADs) and the *in vitro*-characterized 30-nm fiber - remain much more limited. A major contributor to this lacuna is the reliance on short-read sequencing, which limits direct inference of large-scale chromosomal architecture. CheC-PLS promises to help fill this gap.

Our characterization of the meiotic chromosome axis indicates that CheC-PLS correctly captures chromosome-associated sites. Multiple lines of evidence support this assertion. First, we see an accumulation of methylation with prolonged expression. Second, methylation patterns are similar between biological replicates and between meiotic time points. Third, methylation patterns mostly correlate with the results of ChIP-seq experiments. Fourth, methylation accumulates at centromeres, as known for cohesins. Fifth, methylation patterns are distinct from methylation patterns of naked DNA, and do not correlate with sequencing depth or GATC density, arguing against technical artifacts.

An important unknown in the design of CheC-PLS was the *in vivo* kinetics of DNA methylation by dam. This has important implications since very efficient methylation might have introduced background due to methylation by unbound proteins. Very inefficient methylation would have prevented robust methylation signatures. While the exact rate of *in vivo* methylation remains unknown, the gradual accumulation of methylation between 3 and 5 hours indicates that methylation by dam occurs over a time scale of tens of minutes. The strong signal we observe suggests that methylation by diffuse proteins, which is expected to be mostly random, remains limited.

Our analysis of the methylated long reads generated by CheC-PLS illuminates two key aspects of cohesin dynamics that would have been challenging to detect using ChIP-seq or Hi-C. First, we observe a consistent positive correlation between methylated sites on the same sequencing read. We hypothesize that this correlation is caused by loop extrusion, leading to extensive translocation of cohesin along the same DNA molecules, methylating GATC sites along its path. This is supported by the following observations: (1) Correlation does not dramatically diminish with distance (up to 40kb), arguing it is not a result of the passive sliding or Brownian motion of the chromosomes or of the flexible linker between Rec8 and dam. (2) The correlation diminishes upon elimination of Wpl1, which increases the residency time of cohesin on chromosomes and reduces available cohesins to perform loop extrusion. (3) Correlation is eliminated in isolated nuclei, where lack of ATP eliminates loop extrusion. (4) Correlation is lower between highly methylated sites - corresponding to cohesin peaks in ChIP-seq data. These peaks are more stably anchored at the axis and less mobile, resulting in less translocation-mediated correlation.

The second salient feature is the ∼165bp periodicity of short-range correlation. This distance is very close to the 163-175bp preferential distance between nucleosomes in vegetative budding yeast (Chereji et al. 2018). Periodicity is less pronounced in later meiotic time points and is not dramatically affected by the removal of Wpl1. However, it is much stronger in isolated nuclei. We hypothesize that this periodicity stems from the positioning of nucleosomes in uniform orientation relative to cohesins, resulting in preferential methylation of GATC sites on the same position on adjacent nucleosomes, and/or due to preferential methylation of spacers that are also similarly spaced.

The long-reads produces by CheC-PLS can lend unprecedent insight into the organization of genomic regions composed of tandem repeats, including telomeres, centromeres and the rDNA, where short sequencing reads cannot be uniquely mapped. Here we applied CheC-PLS to the rDNA locus, which in budding yeast comprises 100-200 identical 9.1kb tandem repeats. Despite its essential role in ribosome biogenesis and nucleolar organization, and its local effects on recombination (Vader et al. 2011), its native internal organization is poorly characterized (Jiang et al. 2024). We found that cohesins exhibit consistent binding patterns among repeats, and that cohesin exhibit increased occupancy in the rDNA array relative to abutting sequences. Interesting, we find that little evidence that cohesin occupancy is specifically co-regulated between adjacent repeats within the rDNA array or between the repeats and the adjacent non-repeated regions.

In addition to the issues plaguing all ChIP-based approaches, such as perturbative tagging or nonspecific antibodies, the current iteration of CheC-PLS suffers from two specific limitations. The first relates to the reliance on the GATC motif, which limits the resolution to ∼256bp. The effective resolution is likely lower, both due to the uneven distribution of GATC sites and the error rate of methylation calling. Future iterations of CheC-PLS could utilize more promiscuous methyltransferases, such as Hia5 and EcoGII or the cytosine methyltransferases SssI and CviPI (Altemose et al. 2022; X. Yue et al. 2022; Shipony et al. 2020). Denser methylation signal would enable the smoothing of the methylation plots, increasing the confidence in identifying methylated regions at the expense of resolution. Different methyltransferases could also overcome sequence biases in genomic regions of interest (such as the G-rich repeats constituting the telomeres) and allow adaptation of CheC-PLS to organisms with different native methylation patterns.

The second limitation is the flexibility in inducing methyltransferase activity. In the current work, we relied on the native transcriptional pattern of the meiosis-specific Rec8 to express the tethered methyltransferase. This limited our ability to conclusively deduce the patterns of cohesin association in late meiotic time points. Accumulation of methylation also limited the dynamic range of CheC-PLS, dampening the signal at later time points. The ability to deploy CheC-PLS on isolated nuclei could mitigate this issue, and also obviates the need for genome engineering and controls for the potential artifactual effects of methylation. Nonetheless, as our data shows, methylation on isolated nuclei does not fully recapitulate *in vivo* methylation. Notably, methylation at centromeres was affected, and the ability to study dynamic processes was also curtailed.

CheC-PLS offers unique advantages that build on existing genomic approaches, including widely applied approaches such as ChIP-seq and Hi-C, as well as more recently developed approaches that rely on long-read sequencing such as Pore-C and DiMeLo-seq. CheC-PLS adds the ability to study dynamic events and to probe the organization of genomic regions composed of highly repetitive sequences. Its future application to biological processes in diverse model organisms and cell lines promises to shed light on poorly understood features of genome organization.

## Materials and Methods

### Yeast strains

All *Saccharomyces cerevisiae* strains are derivatives of SK1. Methyltransferase gene sequences were inserted in-frame at the 3’ ends of genes at the endogenous loci using recombination-mediated construction. Detailed information on all strains is provided in Supplemental Table 1. Tetrad dissection was performed on Nikon Eclipse Ci microscope with tetrad dissection attachment.

### Yeast cells preparation

Meiosis was induced essentially as described (Brar et al. 2012). Frozen stocks were streaked onto a YPAG (1% Yeast extract, 2% Peptone, 0.01% Adenine hemisulfate, 2% Glycerol) plate for overnight growth to ensure respiration competence. Subsequently, yeast cells were transferred from the YPAG plate to a YPAD (1% Yeast extract, 2% Peptone, 0.01% Adenine hemisulfate, 2% Glucose) plate and incubated for 12 hours. Afterward, cells were transferred to YPAD liquid medium and allowed to grow for 24 hours, harvested and washed twice with water. The washed cells were transferred to BYTA (1% Yeast extract, 2% Bactotryptone, 1% Potassium acetate, 50mM Potassium phthalate) liquid medium and incubated overnight. Following this incubation, the cells were again harvested, washed twice with water, and then transferred to SPO (0.3% Potassium acetate, 0.02% Raffinose) medium at a concentration of 1.85 OD, for induction into meiosis. Cells were incubated in SPO medium for 3-6 hours, shaked at 250 rpm in a flask >×10 volume for proper aeration. Throughout, yeast cells were grown at 30°C.

### High molecular weight DNA extraction

High molecular weight DNA extraction was performed similarly to (Erwan Denis, Sophie Sanchez, Barbara Mairey, Odette Beluche, Corinne Cruaud, Arnaud Lemainque, Patrick Wincker, Valérie Barbe 2018). 1×10^9^ meiocytes were washed with 10 ml of K-sorb (0.1 M KHPO_4_ and 1.2M sorbitol, pH = 6.5) twice. Subsequently, the cells were resuspended in 5 ml of K-sorb, and 50 µl of zymolase (USBiological, Z1004 Zymolyase 100T) and 10 µl of β-mercaptoethanol were added to remove the cell wall. The cell suspension was incubated at 30°C for 40 minutes, with gentle inversion every 15 minutes. The spheroplasts were washed twice with K-sorb (2,000 rpm, 2 minutes), transferred to an Eppendorf tube, and resuspended in TLB buffer (10mM Tris-Hcl, 25mM EDTA, 0.5 w/v SDS). RNAse was added at 1:500 concentration. The cell suspension was incubated at 37°C for 1 hour.

Subsequently, 5 µl of proteinase K was added, and the mixture was incubated at 50°C for 1 hour. The cells were then centrifuged at maximum speed for 1 minute. The supernatant was poured into the phase-lock tubes (Quanta bio, Cat# 2302820), and an equal volume of 25:24:1 phenol:chloroform:isoamyl alcohol was added. The tubes were gently rotated on a nutator for 10 minutes, followed by centrifugation at maximum speed for 10 minutes. The supernatant was transferred to a new phase-lock tube, and the process was repeated. The aqueous phase was collected into a 50 ml tube, and 400 µl of 5M ammonium acetate and 3 ml of ice-cold 100% ethanol were added. Clusters of DNA threads were fished with a pipette and moved into a tube containing 70% ethanol, and then transferred to an Eppendorf tube containing 70% ethanol. After gentle centrifugation (300 rpm) to remove excess ethanol, the DNA was dried at room temperature. Finally, 100 µl of EB buffer or water was added to rehydrate the genomic DNA.

### Nanopore library preparation and sequencing

For nanopore sequencing, we used the RAD004, LSK109 or RBK004 kits (Oxford Nanopore) to maximize the fraction of long DNA reads. The protocol was executed according to the manufacturer’s documentation. Sequencing was conducted using an Oxford Nanopore MinION sequencer, equipped with v9.4 flow cells (ON FLO-MIN106.1), and operated with the MinKNOW software (version 21.02.1).

### Base-calling and methylation calling

Raw nanopore sequencing reads (fast5 files) were base-called using Guppy (Oxford Nanopore Technologies). We further employed minimap2 (H. Li 2018), bwa (H. Li and Durbin 2009), samtools (H. Li et al. 2009), and nanopolish (Simpson et al. 2017) to align the reads to the reference genome of SK1 strain (J.-X. Yue et al. 2017), index the reads in the bam file, and generate eventalign data. We explored two algorithms for detecting adenine methylation, mCaller and Remora (https://github.com/al-mcintyre/mCaller; https://github.com/nanoporetech/remora). mCaller utilizes a statistical approach to detect deviations from the expected current as DNA passes through the sequencing pore (McIntyre et al. 2019). Remora (Oxford Nanopore Technologies) employs deep learning, where a neural network model is trained to recognize methylation patterns using a dataset where the ground truth of methylation is known. To train Remora, we used two *E. coli* strains: *dam^-^ dcm^-^* with no adenine methylation, and a wild-type (*dam^+^ dcm^+^*) strain where essentially all GATC sites are methylated. Both algorithms assign each GATC site a methylation value ranging from 0 to 1, with higher values indicating a greater likelihood of methylation. Our evaluation indicated that Remora performed better on our datasets. Remora demonstrated higher accuracy than mCaller, with lower rates of both false-negative and false-positive calls (Fig. 1e; we used 50% of the sequencing reads to train Remora, and the rest for testing). For all of the analysis below we used a threshold of 0.61, which resulted in a false identification rate of less than 15%, compared with 30% error rate when using mCaller (Fig. 1f).

### Pulse-field gels

DNA was subjected to pulse-field gel electrophoresis (PFGE) as described in (Rog et al. 2009). DNA was separated using CHEF-DR II (Bio-Rad). The DNA ladder used was the CHEF DNA Size Marker (Bio-Rad, Cat# 170-3605).

### Bacterial DNA strains and preparation

All plasmids utilized in this study are detailed in Supplemental File 1. To construct Rec8-dam, Gibson Assembly was employed, using pSB2065 plasmid as the backbone. Plasmids were transfected into *E. coli* strain TH16833, which lacks *dam* and *dcm* genes, serving as a storage host. Strains RP900 (*dam+ dcm+*) and RP8612 (*dam-dcm-*) were used as controls for completely methylated and unmethylated genomes.

### Analysis of ChIP

Rec8 ChIP-seq data, using rabbit Rec8 antiserum and Protein A agarose beads, was downloaded from NCBI (Fajish et al. 2024). This pre-processed data provided relative enrichment on each genome position. To compare this data to the aggregated CheC-PLS data, we used the same window size and step length to analyze the data.

### Statistical analysis

Statistical analyses were conducted using Python’s SciPy library (Virtanen et al. 2020). Specifically, Pearson correlation coefficients (P corr) were calculated using the pearsonr function from scipy.stats. Two-sample t-test were implemented by the ttest_ind function from scipy.stats. P-values less than 0.05 were considered statistically significant.

### Generating shuffled datasets

To create shuffled datasets, all reads spanning each GATC site were identified. We compiled all methylation values from these reads, randomized them, and then reassigned them. As a result, the average methylation at each GATC site, as well as the distribution of read lengths and genomic coverage were identical to the CheC-PLS data, although it lacked any correlation between methylation sites. The methodology for generating this simulated data is detailed and available on GitHub.

### Cross-correlation analysis

To examine CoC between two GATC sites, we analyzed all reads spanning both sites for their methylation status. For each site, the methylation status was set to either 1 (methylated) or 0 (unmethylated), based on the 0.61 threshold. This resulted in four possible scenarios: both sites unmethylated (0,0), left site methylated and right site unmethylated (1,0), left site unmethylated and right site methylated (0,1), and both sites methylated (1,1). We quantified the fraction of reads corresponding to each scenario. These fractions were then used to calculate the Correlation Coefficient (CoC), as described by the following equation: ln(CoC) = ln(*Q / (M*N)).* Q = P(1,1), M = P(1,0)+P(1,1), N= P(0,1)+P(1,1) (Zhang et al. 2014).

In all experiments we observed a minor but distinct population of reads that lacked methylation, presumably due to failure to enter the meiotic cell cycle. For ln(CoC) analyses we excluded reads that harbored less than 4.8% methylation reads (Supplementary Fig. 1h).

### rDNA assembly & analysis

We utilized the publicly available sequence of budding yeast chromosome XII as a backbone. This genome contained two rDNA repeats, to which we manually added 18 identical repeats to create a genome containing 20 repeats. We used this modified genome to align all reads proximal to the rDNA locus and assess their methylation status within the rDNA locus. The rest of the analysis was performed as above.

### dam methylation of naked DNA

dam enzyme (NEB, Cat# M0222S) was used according to the manufacturer’s instructions. 5 ug of genomic DNA from wildtype yeast strain was incubated with dam for 1 hour at 37°C in a buffer containing 80 µM S-adenosylmethionine (SAM), cleaned up using 25:24:1 phenol:chloroform:isoamyl alcohol, and sequenced and analyzed as above.

### CheC-PLS on isolated nuclei

GBP-dam was cloned and purified by GenScript. The GBP sequence (FROM WHERE?) was fused to the dam sequence (FROM WHERE?) with three v5 linkers. Meiotic nuclei were obtained by synchronizing Rec8-GFP strain to undergo meiosis, as described above. After 4 hours in SPO, nuclei were isolated according to the (Greenwood et al. 2018). Successful isolation of nuclei was determined by micrococcal nuclease (1 ul, 300 units) digestion, which yielded nucleosome-sized bands. Approximately 3.6×10^7^ isolated nuclei were incubated with 8 ug of GBP-dam for one hour, in conditions similar to those used for dam methylation of naked DNA. Genomic DNA isolation, sequencing and methylation calling were conducted as above.

### Authors contributions

All experiments and analysis were performed by KX, except the nuclei isolation experiments that were carried out by YZ. JBB contributed to code development. TAS contributed sequencing information for earlier iterations of CheC-PLS. NP provided mentoring. MPM participated in experimental design and mentoring. KX and OR conceptualized the project, designed the experiments and wrote and edited the paper.

## Acknowledgements

All members of the Rog lab for discussions; scientific illustrator Maria Diaz de la Loza for graphical work; Colin Dale and Li Szhen (Michelle) Teh for the use of PFGE machine; Peter Ames and Sandy Parkinson for *E. coli* strains; Jian Huang for generating the ColabFold projection image for cohesin and tagged dam; Luke Berkowitz for yeast strains; and Joe Carrier, Emily Parnell, Lexy von Diezmann, Jamie Gagnon, Michael Werner, Mike Shapiro, Julie Cooper, Aaron Quinlan, Richard Clark and Aaron Fleming for discussions and advice. This work is supported by NIGMS grants R35GM142749 (to MPM) and R35GM128804 (to OR).

