## Supplemental Figures for "Decoding chromosome organization using CheC-PLS: chromosome conformation by proximity labeling and long-read sequencing"

### Supplementary Figure 1

**a**

| Strains | Heterozygous Viability | Haploid Viability | Homozygous Viability | Sporulation | Spore Viability | Methylation detected |
|---|---|---|---|---|---|---|
| *mer2-M.SssI* | √ | √ | √ | √ | × | × |
| *red1-M.SssI* | √ | √ | √ | × | | × |
| *mer2::M.CviPI* | √ | √ | √ | √ | × | x |
| *mer2-dam* | √ | √ | √ | √ | × | |
| *hop1-dam* | √ | √ | √ | √ | × | |
| *rec8-dam* | √ | √ | √ | √ | √ | √ |
| *rec8-M.CviPI* | √ | √ | √ | × | | |
| *rec8-EcoGII* | √ | √ | √ | × | | × |

**c**

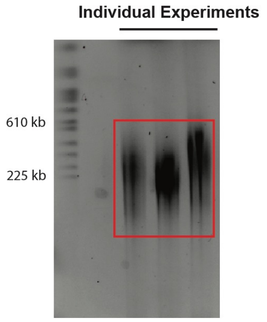


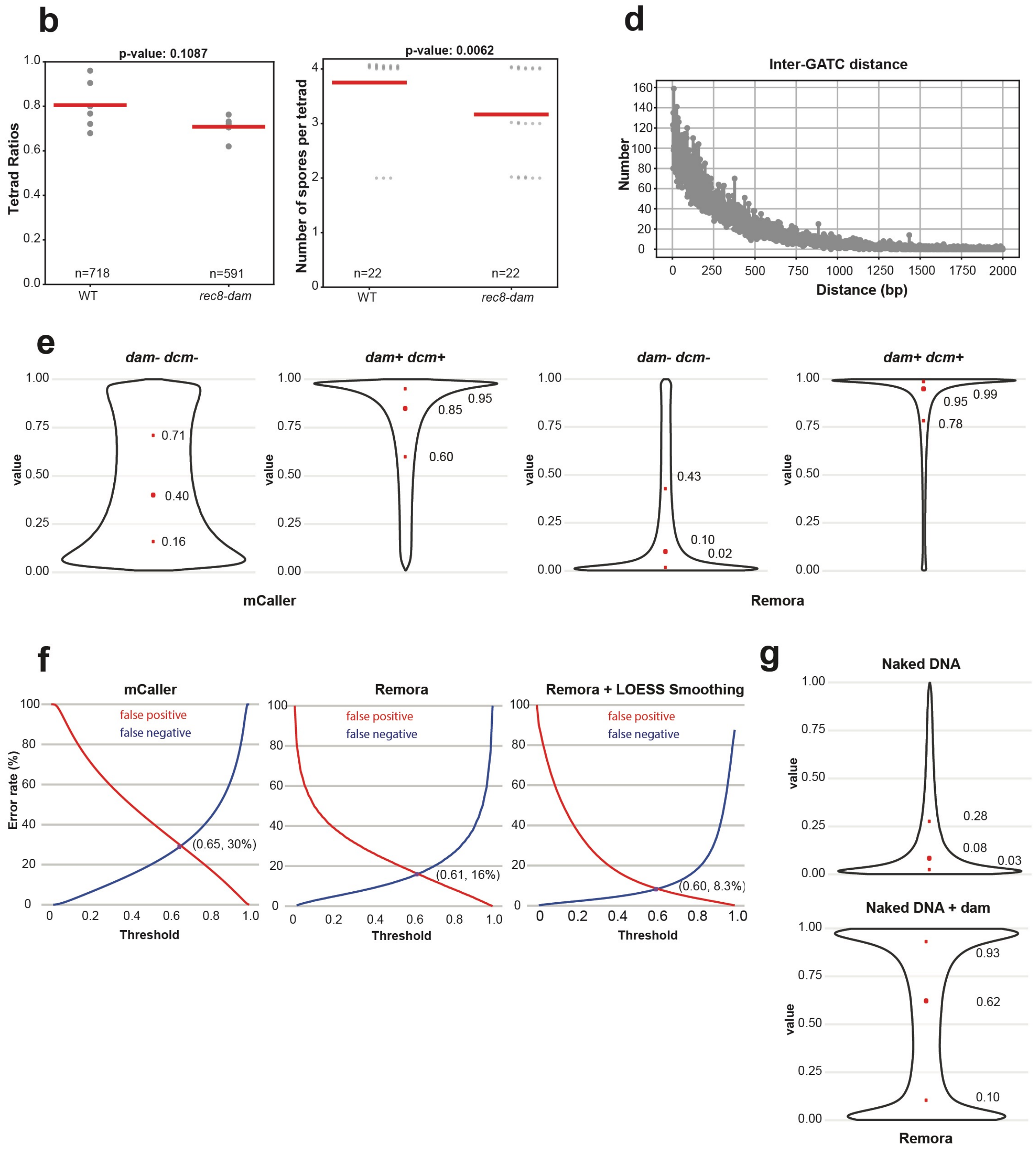


7

#### Supplementary Figure 1

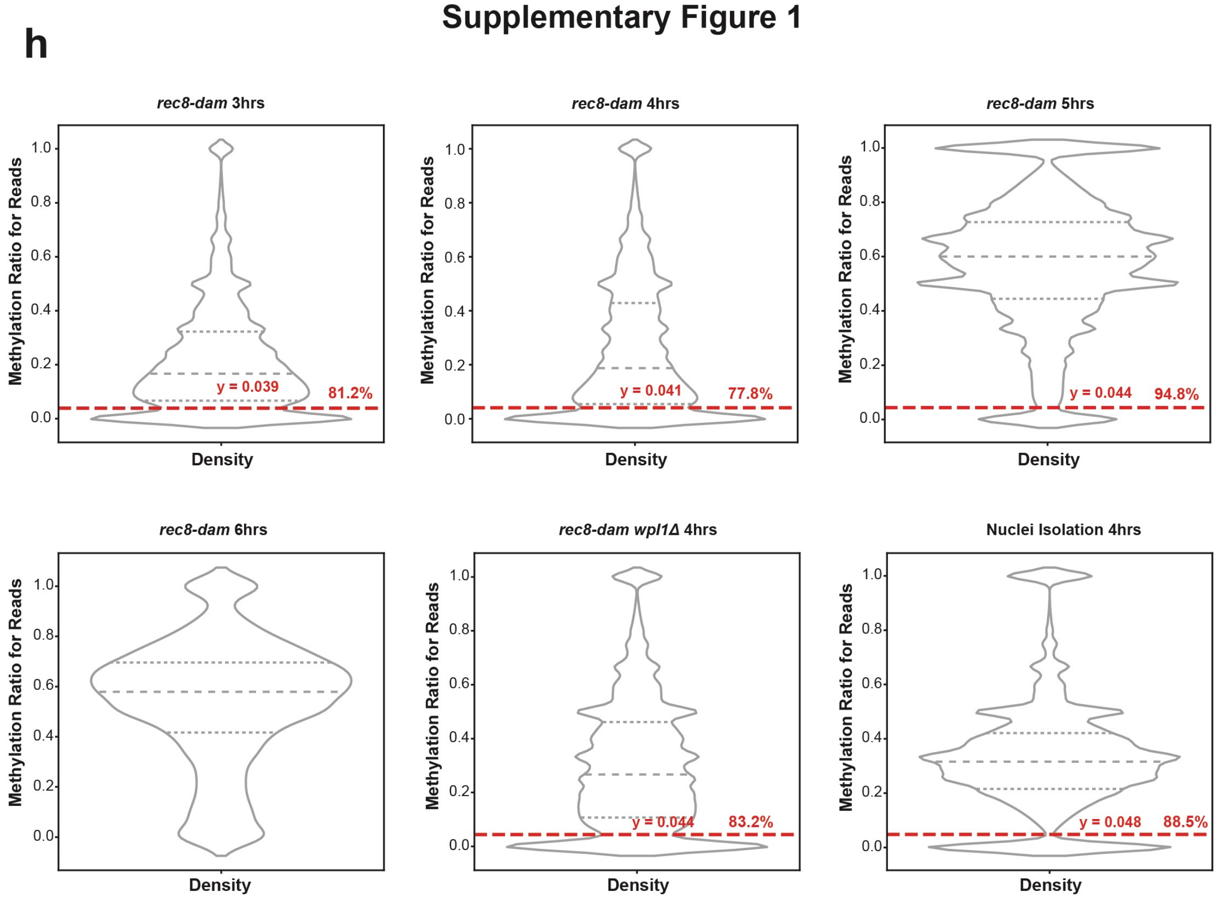


**Supplementary Figure 1. Tagging Rec8 with dam**

(a) Functional characterization of various meiotic factors tethered to DNA methyltransferases. Except *rec8-dam*, all other fusion proteins exhibited perturbed meiotic functions. (b) *rec8-dam* forms tetrads in similar rates to wild-type yeast, although slightly fewer tetrads contain four viable spores. (c) High molecular weight DNA was extracted from three individual experiments and then subjected to pulsed-field gel electrophoresis. The shorter read lengths observed in the CheC-PLS sequencing output, compared with the results from the pulsed-field gels, suggest that shearing occurred during library preparation. (d) Dot plot showing the genomewide distribution of GATC sites by distance. We did not find evidence for the 165bp periodicity, or any other distinct pattern. (e) Violin plots of total methylations calls by mCaller and Remora on *dam- dcm-* (negative control) and *dam+ dcm+* (positive control) *E. coli* strains. Red dots indicate 25th, 50th and 75th percentiles. (f) False negative and false positive rates for different threshold values, based on data from panel e. A threshold of 0.61 for methylation detection using Remora yielded the lowest false rate. LOESS smoothing can improve methylation calling at the expense of resolution. (g) Violin plots of total methylations calls on naked budding yeast genomic DNA treated, which lacks endogenous adenine methylation, and naked DNA incubated with recombinant dam enzyme *in vitro*. Red dots indicate 25th, 50th and 75th percentiles. (h) Violin plots of average methylation per sequencing read from six different experiments. The red dashed lines indicate the bottleneck in each experiment, which likely represent DNA from cells that fail to be induced into meiosis. Reads below this threshold (4.8%) were removed from the analysis. Red percentages indicate the fraction of reads included in the CoC analysis.

### Supplementary Figure 2

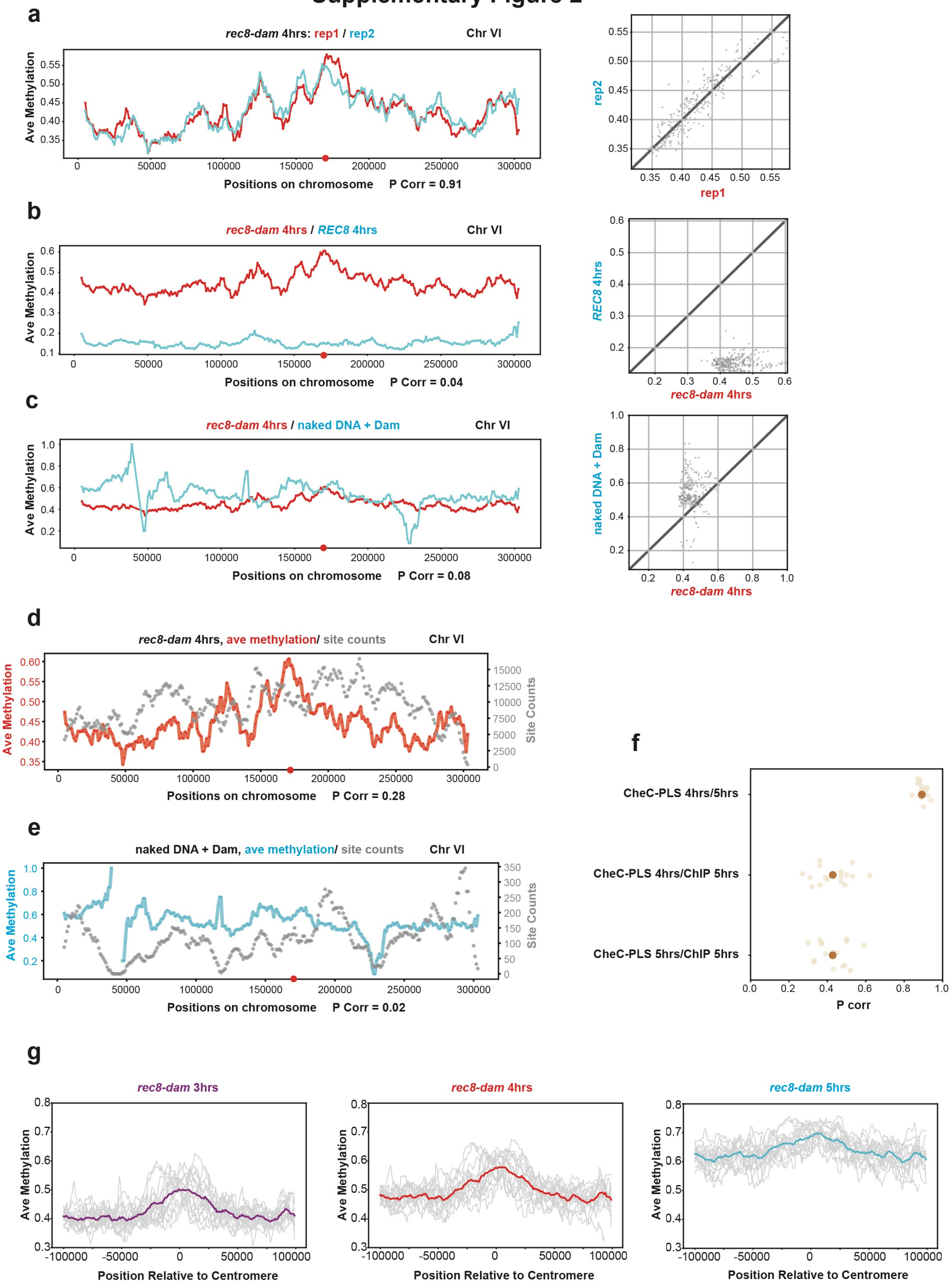

**Supplementary Figure 2. Comparison of running methylation plots on chromosome VI under different conditions**

(a) Averaged methylation for two biological replicates. The window size is 10 kb. The red dot indicates the centromere. Right, scatter plot between the two replicates. (b) Averaged methylation for *rec8-dam* and untagged *REC8* strain. The window size is 10 kb. The red dot indicates the centromere. Right, scatter plot between the two strains. (c) Averaged methylation across *rec8-dam* and naked DNA incubated with dam enzyme. The window size is 10 kb. The red dot indicates the centromere. Right, scatter plot between the two experiments. (d) Averaged methylation of *rec8-dam* and the GATC site count - the total number of GATC sites from all the reads within the 10kb window. The red dot indicates the centromere. (e) Averaged methylation of naked DNA with dam incubation and the GATC site count. The window size is 10 kb. The red dot indicates the centromere. (f) Averages of Pearson correlation for all 16 chromosomes under different conditions. The dark yellow dots represent the average. (g) Averaged methylation around the centromeres for all 16 chromosomes (grey lines) at three time points. The window size is 10 kb.

### Supplementary Figure 3

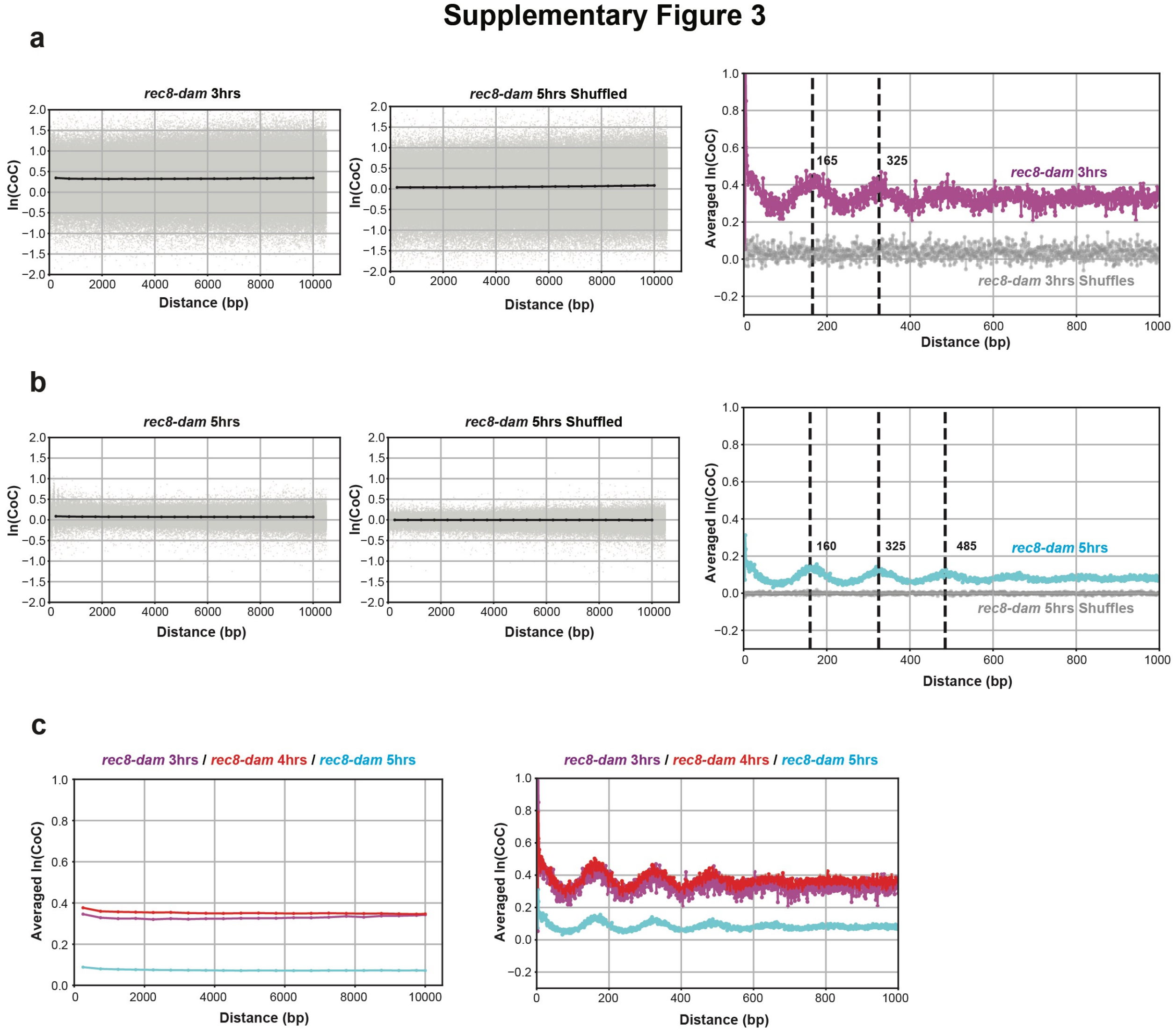

#### Supplementary Figure 3

d

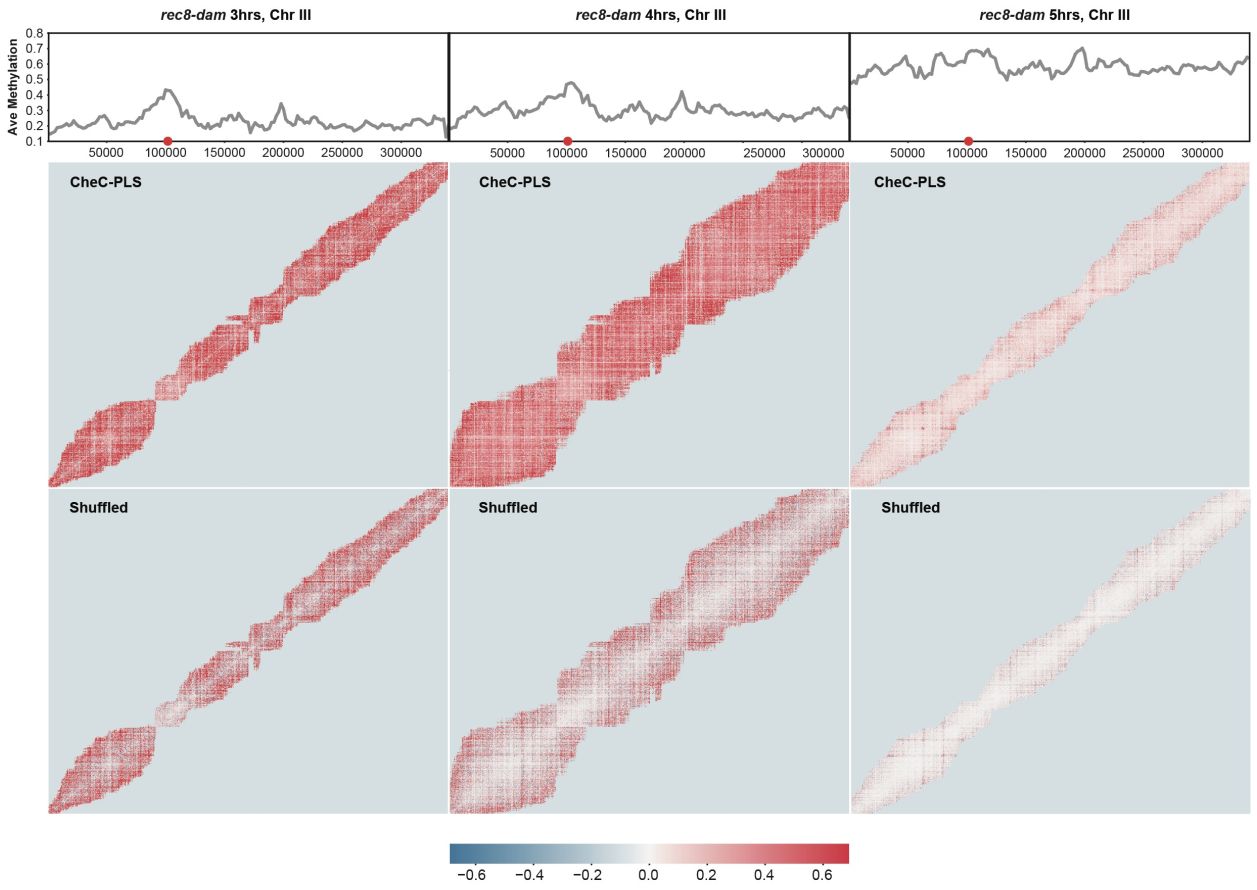


**Supplementary Figure 3. ln(CoC) by distance indicating Rec8 sliding on chromosomes at different time points after induction into meiosis.**

(a) Averaged ln(CoC) by distance between each pair of sites in CheC-PLS *rec8-dam* 3 hours and shuffled data. Grey dots indicate each pair of sites, and black dots indicate the averaged ln(CoC). Right, Zoomed-in view with no binning. The CheC-PLS *rec8-dam* 3 hours is shown in magenta and the shuffled dataset in grey. (b) Averaged ln(CoC) by distance between each pair of sites in CheC-PLS *rec8-dam* 5 hours and shuffled data. Grey dots indicate each pair of sites, and black dots indicate the averaged ln(CoC). Right, Zoomed-in view with no binning. The CheC-PLS *rec8-dam* 5 hours is shown in cyan and the shuffled dataset in grey. (c) Comparison of averaged ln(CoC) by distance between *rec8-dam* 3 hours (magenta), 4 hour (red) and 5 hours (cyan). The analysis spans from 0 to 10 kb with a bin size of 500 bp (left) and from 0 to 1 kb without binning (right). (d) Heatmap of ln(CoC) for each pair of sites across chromosome III for *rec8-dam* at 3, 4 and 5 hours. ln(CoC) ranges from blue to red. Top, average methylation plot along the chromosome.

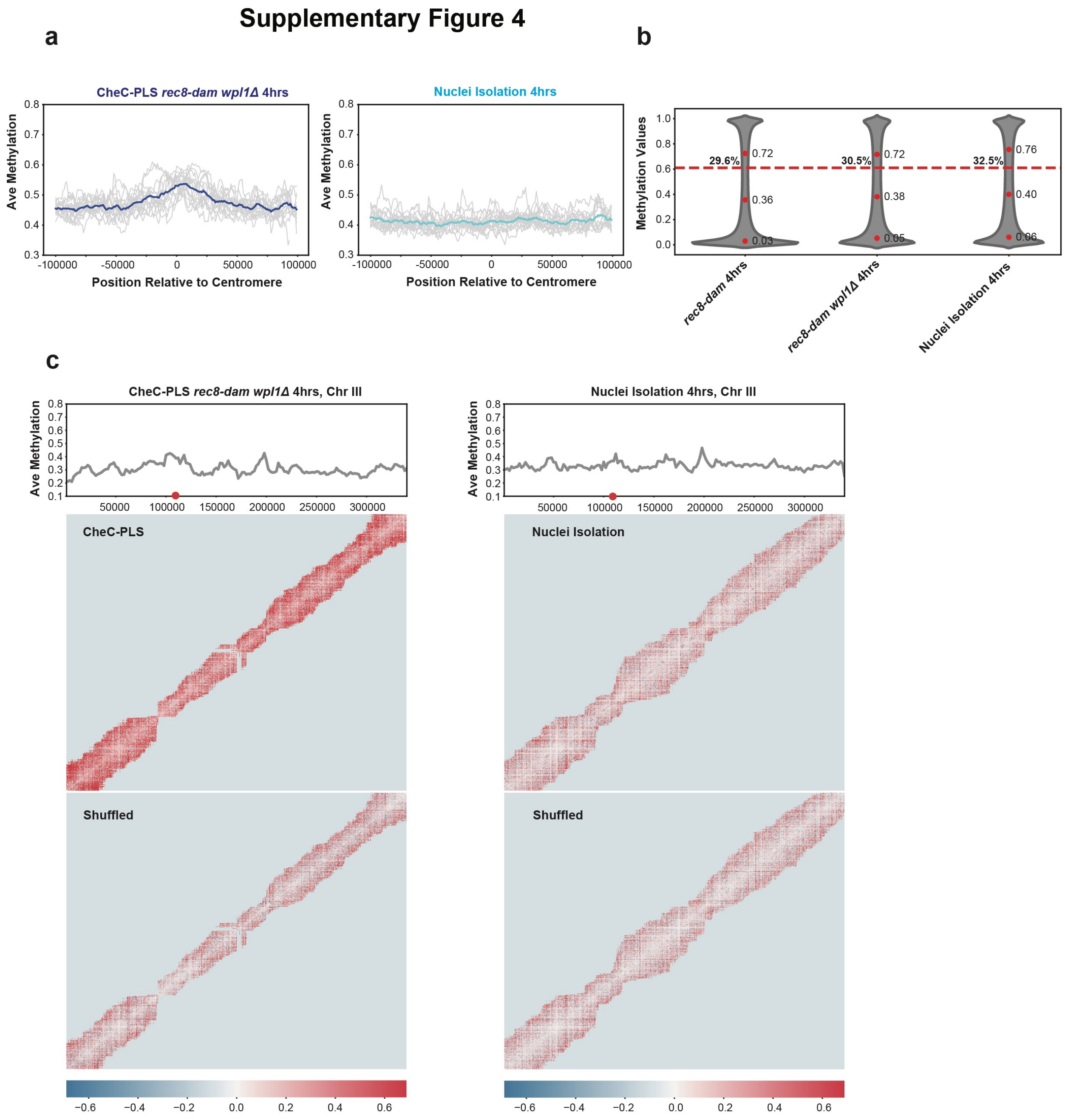


**Supplementary Figure 4. Shared and distinct features of *wpl1Δ* and nuclei isolation compared to CheC-PLS.**

(a) Averaged methylation around the centromeres for all 16 chromosomes (grey lines) in *rec8-dam wpl1Δ* at 4 hours and nuclei isolation at 4 hours. The window size is 10 kb. (b) Violin plots of total methylation calls and the fraction of reads with methylation rates above 0.61 for each experiment (dashed lines). The red dots indicate values for 25th, 50th and 75th percentiles. (c) Heatmap of ln(CoC) for each pair of sites across chromosome III. ln(CoC) ranges from blue to red. Top, average methylation plot along the chromosome.

#### Supplementary Figure 5

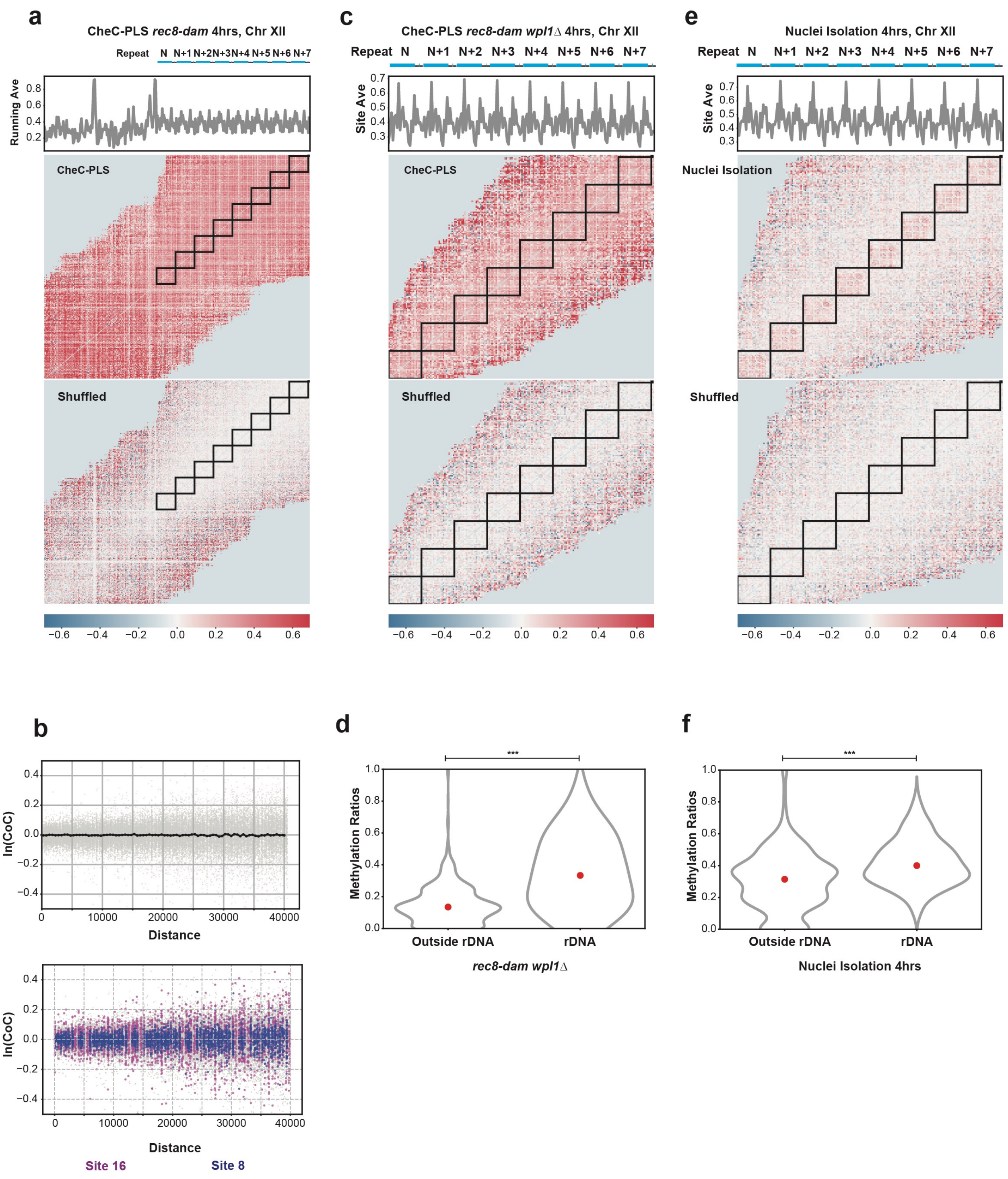


**Supplementary Figure 5. Higher-order chromosome structure in the rDNA region in WT and *wpl1Δ* cells, with distinct patterns observed using the nuclei isolation method.**

(a) Heatmap of ln(CoC) in pooled reads of the rDNA region, showing the region right before rDNA and eight rDNA repeats in CheC-PLS *rec8-dam* 4 hours. Top, average methylation plot along this region. The window is 10 kb, with a step size of 1 kb. (b) Top, ln(CoC) by distance in pooled reads mapping to the rDNA in shuffled data. Bottom, site 16 (magenta) are no longer associated with high ln(CoC), while site 8 (blue) are no longer associated with low ln(CoC) in the shuffled data. (c) Heatmap of ln(CoC) in pooled reads of the rDNA region from repeat N to repeat N+7 in *rec8-dam wpl1Δ* cells, showing inter- and intra-repeat correlations. (d) A violin plot showing the average methylation levels in 1000 randomly selected 9.1 kb windows outside the rDNA and averaged methylation levels for all rDNA repeats in *rec8-dam wpl1Δ* cells. (e) Heatmap of ln(CoC) in pooled reads of the rDNA region from repeat N to repeat N+7 in nuclei isolation 4 hours, showing intra-repeat correlations. (f) A violin plot showing the average methylation levels in 1000 randomly selected 9.1 kb windows outside the rDNA and averaged methylation levels for all rDNA repeats using the nuclei isolation process.
